# A causal role of the human left temporoparietal junction in computing social influence during goal-directed learning

**DOI:** 10.1101/2022.06.13.495824

**Authors:** Lei Zhang, Farid I. Kandil, Ke Zhao, Xiaolan Fu, Claus Lamm, Claus C. Hilgetag, Jan Gläscher

## Abstract

The human temporoparietal junction (TPJ) is a brain area crucial for processing social information. Although brain stimulation studies have started to explore the causal function of TPJ under social contexts, few have explicitly considered bilateral TPJ as target regions. Here, leveraging non-invasive continuous theta-burst stimulation (cTBS) and hierarchical Bayesian computational modeling, we tested whether left or right TPJ (with vertex as control) is causally involved in how dissenting choices by others influence individuals’ choice adjustments in goal-directed learning. In our social learning paradigm, participants (N = 31) first made their private decision, and then were allowed to re-adjust their choices after observing choices of four other players. Behaviorally, we show that disruption of the left, but not the right TPJ, weakened participants’ choice adjustment and delayed their response speed when confronted with dissenting information from the other players. Computationally, disrupting activity in the left TPJ attenuated the degree of computing social influence during choice adjustment, whereas the extent to which how observational learning from others’ choices was integrated into direct learning remained intact. Together, our results provide evidence for the causal role of left TPJ in social influence during goal-directed learning and shed light on the relational function (with respect to oneself) of the TPJ in social cognition.

## INTRODUCTION

The human temporoparietal junction (TPJ) is a brain area crucial for processing social interaction, Theory of Mind (ToM), and self-other distinction (Ruff, Fehr, 2014; Schaafsma et al., 2015; Deschrijver and Palmer, 2020; Rusch et al., 2020; Schurz et al., 2021). Despite a recent focus on the right TPJ as a primary site for representing such social information, many functional magnetic resonance imaging (fMRI) studies have indeed reported bilateral activation of TPJ in social influence in goal-directed learning (Zhang and Gläscher, 2020), strategic social decision-making (Konovalov et al., 2021), and goal emulation in observational learning (Charpentier et al., 2020), and these results are consistent with large-scale, automated meta-analytic findings reported on NeuroSynth (Yarkoni et al., 2011). For example, we have previously documented the neurocomputational role of bilateral TPJ in computing social influence in goal-directed learning, in which participants were able to re-adjust their choices after observing choices of four other players in the same learning environment (Zhang and Gläscher, 2020). To establish the causal relationship between TPJ and social information processing, several brain stimulation studies have demonstrated a causal involvement of right TPJ in moral judgments (Obeso et al., 2018), social norm violation of others (Baumgartner et al., 2014), strategic social interaction (Hill et al., 2017), and self-other distinctions (Bukowski et al., 2020). However, most of these studies have, in fact, only stimulated right TPJ, and not explicitly tested the differential contributions of left and right TPJ to these phenomena.

To bridge this gap, here we employed a modified version of our social learning task and explicitly tested whether left or right TPJ was casually involved in representing dissenting social information when individuals were about to make choice adjustments. With the well-established continuous theta-burst stimulation (cTBS) protocol (Huang et al., 2005) to left and right TPJ and hierarchical Bayesian reinforcement learning models (Zhang et al., 2020), we reported that left, but not right, TPJ was causally related to the computation of social influence during goal-directed learning.

## RESULTS

Human participants (N = 31, 17 females) performed a modified version of our social learning task (Zhang and Gläscher, 2020), which relies on a probabilistic reversal learning paradigm (**Figure 1A**; Gläscher et al., 2009). On each trial, participants made their initial choice (Choice 1), and after being informed about choices from the other four players, participants were allowed to re-adjust their final choice (Choice 2), before receiving the outcome. Participants were incentivized to earn as much money as possible. As we demonstrated previously (Zhang and Gläscher, 2020), the multiple reversal structure (**Figure 1B**) entailed adequate uncertainty such that it was much wiser for the participants to consider the other players’ decisions for better detection of changes after each reversal. Crucially, participants were informed that the other four players were intelligent computer algorithms (see **Materials and Methods**), which were able to learn from reward feedback as well as learn from others’ choices – including the choices of the participant. The algorithms were in fact simulated from our previous computational model that best matched human behavior in the same task setup (Zhang and Gläscher, 2020). All participants underwent three cTBS sessions in a counterbalanced order (**Figure 1D**). We conducted such a fully randomized within-subject, rather than between-subject, design so as to minimize heterogeneity between different groups, thus maximizing the statistical power in estimating cTBS effects (Charness et al., 2012). The stimulation sites included the vertex (as a non-active control region) and right as well as left TPJ, the Montreal Neurological Institute (MNI) coordinates of which (right: [50, –60, 34]; left: [–48, –62, 30]; **Figure 1E**) were extracted based on our previous finding. A neuronavigation pipeline (Polanía et al., 2018) was implemented to obtain stimulation targets tailored to each participant’s brain anatomy.

**Figure 1.**
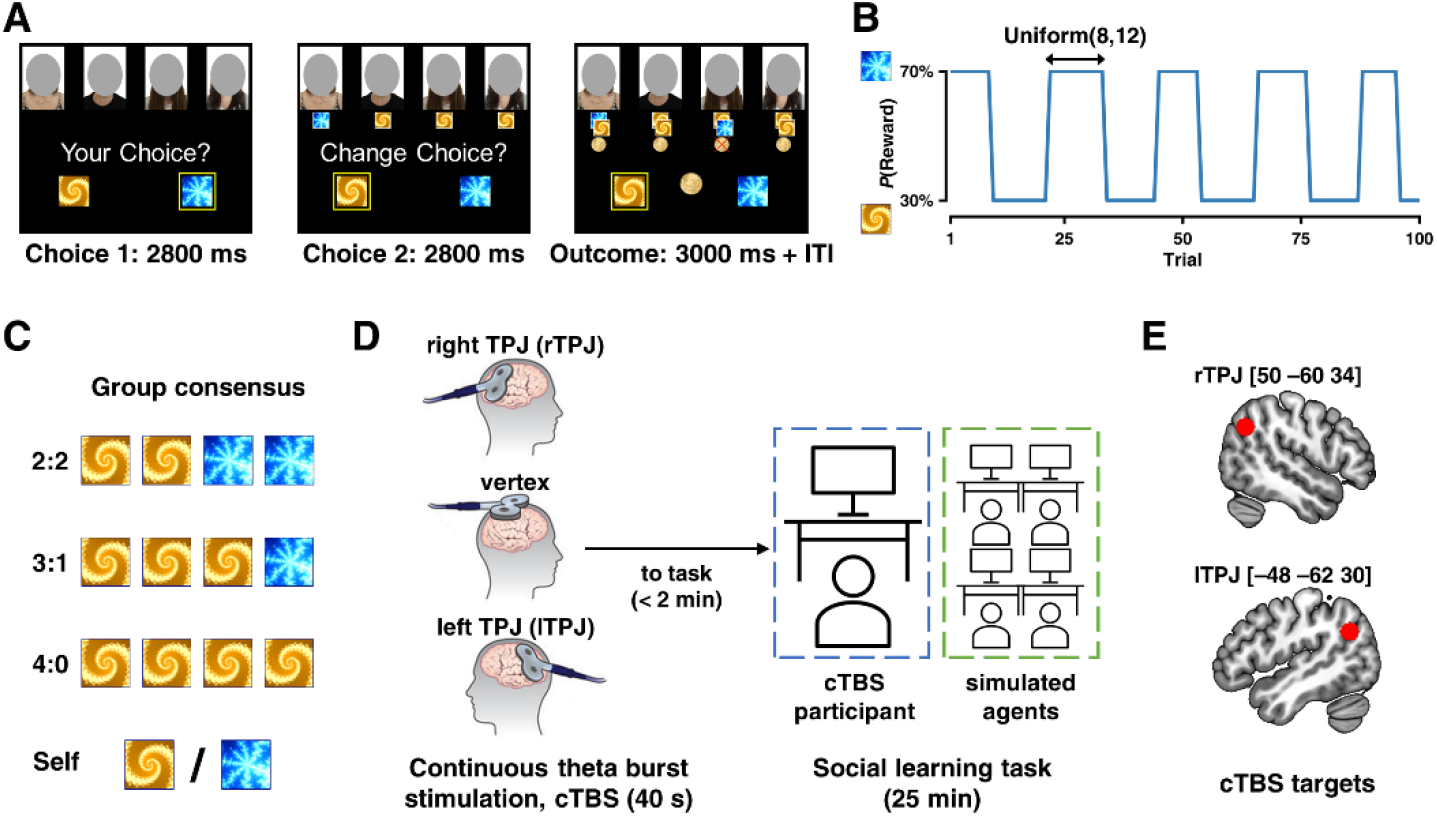
Experimental design. **(A)** Task design. On each trial of the experiment, participants (N = 31) made an initial choice (Choice 1), and after observing the choices from the other four players, participants were asked to make adjustments (Choice 2), followed by the outcome. Three stimulus pairs (one pair shown here) were used with a counterbalanced order across stimulation sites. **(B)** Example reward schedule. Reward contingency reversed every 8–12 trials, following a uniform distribution. **(C)** Illustration of group consensus (perspective from each participant). Note that this illustration was never shown to the participants. **(D, E)** Neurostimulation protocol. After a short practice round, participants received individually localized continuous theta-burst stimulation (cTBS) of areas of interest, right TPJ (Montreal Neurological Institute [MNI] target coordinates [50, –60, 34]) and left TPJ (MNI [–48, –62, 30]), with the vertex as a non-active control region. Participants underwent the experiment in an adjacent testing room next. The order of left or right TPJ cTBS was counterbalanced across participants with vertex stimulation always between the TPJ stimulation to reduce potential interference effects due to hemispheric crosstalk.

### Model-agnostic effects of cTBS on choice behavior

We sought to examine whether stimulating TPJ altered choice switching, response speed, and choice accuracy during participants’ choice adjustment after social influence (i.e., Choice 2). Across all three analyses, we were primarily interested in the stimulation effect on the interaction between the relative direction of the group (with vs. against one’s own first choice) and group consensus (2:2, 3:1, and 4:0, indicated from participants’ perspective; **Figure 1C**), as this interaction was one of the major findings in our previous study (Zhang and Gläscher, 2020). Thus, we computed the difference between the “against” and “with” conditions at each consensus level as the dependent variable for each measurement of interest, and fitted 3 (cTBS sites: right TPJ, vertex, left TPJ) × 3 (group consensus: 2:2, 3:1, and 4:0) linear mixed models (LMM).

We first assessed whether stimulating TPJ reduced choice switch probability (**Figure 2A**, showing the actual data instead of the difference measurement), namely, how likely participants switched to the other choice alternative after observing the four choices of the other players. The LMM revealed a significant main effect of consensus (F_2,52_ = 11.460, p < 0.001; β_consensus_3:1_ = 0.241, p = 0.007, β being the standardized coefficient; β_consensus_4:0_ = 0.367, p < 0.001), and a significant stimulation × consensus interaction (F_4,180_ = 3.897, p = 0.005). The main effect of stimulation remained insignificant (F_2,30_ = 1.402, p = 0.262). This two-way interaction was largely driven by the difference in the “4:0” condition, such that disrupting activity in the left TPJ significantly reduced choice switch probability relative to the right TPJ (t_64_ = 3.587, p = 0.018, Tukey corrected). No other pairwise comparisons yielded significant results. Relatedly, we asked whether stimulating TPJ affected response time (RT) during participants’ choice adjustment (**Figure 2B**). With the same LMM setup, we found a trend level effect of stimulation (F_2,80_ = 2.400, p = 0.097), and the two other effects were not significant (consensus: F_2,43_ = 0.121, p = 0.887; interaction: F_4,144_ = 1.193, p = 0.316). However, we found a significant LMM effect of interaction (β_lTPJ×consensus_4:0_ = 0.197, p < 0.028), suggesting that the RT was longest when the activity in the left TPJ was disrupted in the consensus 4:0 condition. Last, now that we had some indication that stimulating the left TPJ impacted both choice switch probability and RT, we sought to examine whether the accuracy (i.e., deciding on the more rewarded option) of Choice 2 was also altered (**Figure 2C**), based on that we previously had observed an increased performance after incorporating others’ choices (Zhang and Gläscher, 2020). Following the same LMM setup, we observed a significant main effect of consensus (F_2,130_ = 4.66, p = 0.011; β_consensus_3:1_ = 0.326, p = 0.020). The main effect of stimulation (F_2,34_ = 0.490, p = 0.617) and the interaction (F_4,130_ = 0.524, p = 0.719) were not significant.

**Figure 2.**
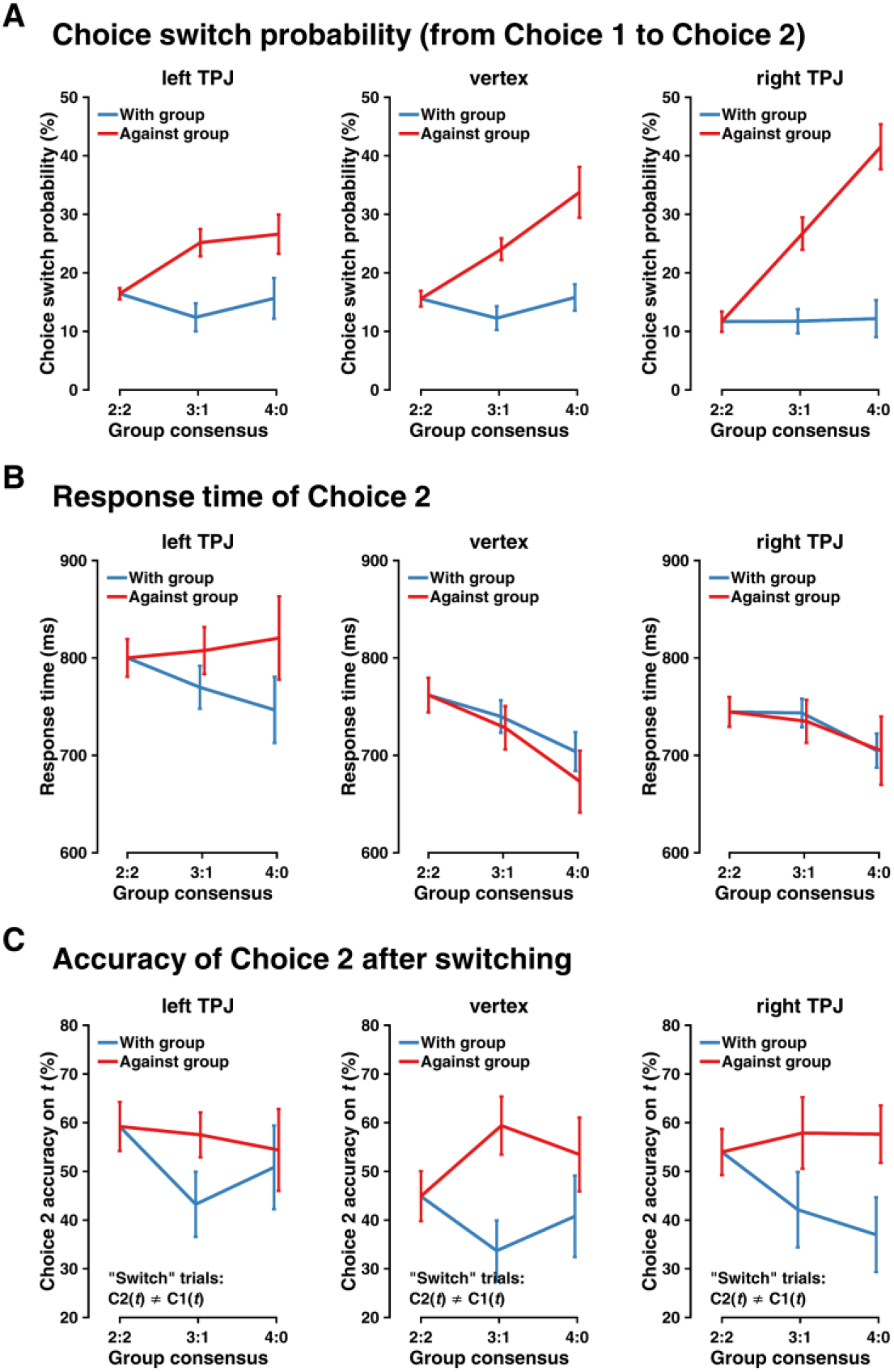
Neurostimulation effects on behavior. **(A)** cTBS effects on participants’ choice switch probability, with a focus on the interaction between group consensus (choice consensus formulated by each computer algorithm; as Figure 1C) and relative direction (with versus against) of the group. Lines indicate means ± within-subject standard error. **(B)** cTBS effects on the response time of participants’ choice switching. Format is as in (A). **(C)** cTBS effects on participants’ choice accuracy after switching. Format is as in (A).

Together, our model-agnostic analyses showed that when the neural activity in the left TPJ was disrupted by cTBS, participants were less inclined to adjust their choices and the associated response speed of making adjustment was remarkably reduced, especially when participants’ choices were contradicted by the others’ choices.

### Computational Mechanisms of social influence in goal-directed learning

Using computational modeling, we aimed to formally quantify latent mechanisms underlying how social influence was computed on a trial-by-trial basis and to uncover nuanced computational contributions to the behavioral differences across cTBS sites. Although we focused on how disruption of the TPJ impacted behavior after receiving choices from the others when outcomes were delivered, participants were also able to learn the others’ choice-outcome combination and integrate such observational learning (through vicarious valuation) into their own valuation on the next trial. We thus constructed our winning model (**Figure 3A**; see **Table 1** for model comparison) following the model development process similar to and detailed in our previous computational account (Zhang and Gläscher, 2020). This procedure subsequently allowed us to investigate whether disruption of TPJ altered vicarious valuation in social learning.

**Figure 3.**
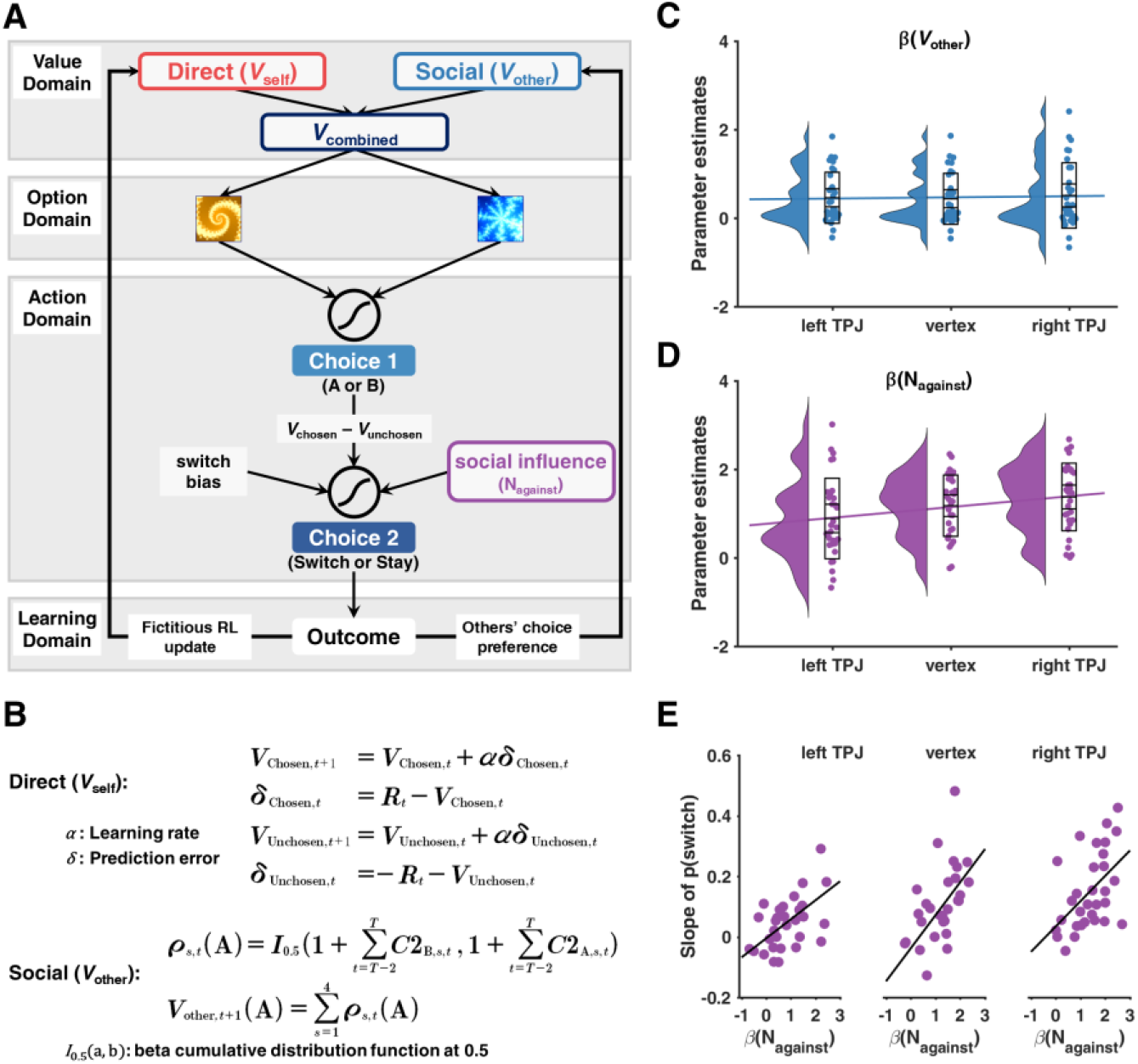
Computational modeling and cTBS effects on model parameters. **(A)** Schematic representation of the winning computational model (M6). Participants’ initial behaviors (Choice 1) were accounted for by value signals updated from both direct learning (*V*_self_) and social learning (*V*_other_); choice adjustments (Choice 2) were ascribed to the valuation of initial behaviors (*V*_chosen,*t*_ – *V*_unchosen,*t*_) and social influence from the other players (N_against_); *V*_self_ was updated via a fictitious reinforcement learning model, while *V*_other_ was updated through tracking other players’ choice preference in past trials. **(B)** Computations of *V*_self_ and *V*_other_. V, value; R, outcome; α, learning rate; δ, reward prediction error; ρ, others’ choice preference; C2_A/B,s_, others’ choices; *I*_0.5(a,b)_: beta cumulative distribution function at 0.5. **(C)** cTBS effects on participants’ degree to integrate *V*_other_ into their valuation (β(*V*_other_)). Violin plots show Kernel density estimation; box plots show mean, standard error, and standard deviation; dots show individual parameter estimates. **(D)** cTBS effects on participants’ computation of social influence (β(N_against_)). Format is as (C). **(E)** cTBS effects on the relationship between β(N_against_) and participants’ susceptibility to social influence (i.e., slope of switch probability calculated from Figure 2A).

**Table 1.** Candidate computational models, model comparison, and winning model’s parameters. # Par., number of free parameters at the individual level per stimulation condition; ΔLOOIC, leave-one-out information criterion relative to the winning model (lower LOOIC value indicates better out-of-sample predictive accuracy); weight, model weight calculated with Bayesian model averaging using Bayesian bootstrap (higher model weight value indicates higher probability of the candidate model to have generated the observed data); HDI, highest density interval estimated from the posterior distributions. M4 (in bold) is the winning model. See Materials and Methods for description of all candidate models.

| Class | Model | Description | # Par. | ΔLOOIC | Weight |
| --- | --- | --- | --- | --- | --- |
| Non-social model<br>(baseline) | M1 | Fictitious RL | 4 | 283 | 0 |
| Social model:<br>social influence | M2 | M1 + social influence | 5 | 117 | 0 |
| Social models:<br>social influence | M3 | **M2 + others' RL update** | 8 | 100 | 0 |
| +<br>social learning | M4 | **M2 + others' action preference** | 6 | – | 0.811 |
|  | M5 | **M2 + others' current reward** | 6 | 35 | 0.189 |
|  | M6 | **M2 + others' cumulative reward** | 7 | 80 | 0 |
| M4 model parameters (group-level mean and 95% HDI) |  |  |  |  |  |
| Parameter | left TPJ | Vertex | right TPJ |  |  |
| $\alpha$ | 0.25 [0.15 0.38] | 0.30 [0.20 0.41] | 0.23 [0.12 0.33] | | |
| $\beta(V_{\text{self}})$ | 1.85 [1.25 2.41] | 1.73 [1.18 2.20] | 1.92 [1.37 2.58] | | |
| $\beta(V_{\text{other}})$ | 0.45 [0.14 0.73] | 0.43 [0.14 0.73] | 0.50 [0.19 0.86] | | |
| $\beta(\text{bias})$ | –2.06 [–2.44<br>–1.67] | –2.17 [–2.52<br>–1.79] | –2.49 [–2.90<br>–2.12] | | |
| $\beta(V_{\text{chosen}} - V_{\text{unchosen}})$ | –0.65 [–0.95<br>–0.38] | –0.56 [–0.81<br>–0.29] | –0.76 [–1.04<br>–0.48] | | |
| $\beta(N_{\text{against}})$ | 0.87 [0.37 1.41] | 1.01 [0.55 1.49] | 1.50 [0.95 2.02] | | |

On each trial, the option value of Choice 1 (A or B) was modeled as a linear combination between values from direct learning (*V*_self_) and values from social learning (*V*_other_). After observing choices from the others, participants’ Choice 2 (switch or stay) was modeled as a function of two counteracting influences: (a) the group dissension (N_against_) representing the social influence, and the value difference between participants’ chosen and unchosen options (*V*_chosen,C1,t_ – *V*_unchosen,C1,t_), representing the distinctiveness of the current value estimates. Lastly, when all outcomes were delivered, *V*_self_ was updated using the fictitious update reinforcement learning (RL) model (Gläscher et al., 2009; Zhang and Gläscher, 2020), whereas *V*_other_ was updated through tracking the other four players’ choice preference history (i.e., the others’ decisions in the recent past; **Figure 3B**). Considering the within-subject structure in our experimental design, we implemented the within-subject effect coding scheme with explicit variance-covariance matrices (see Star Methods) to more accurately reflect the interdependency across conditions for each participant under the hierarchical Bayesian analysis framework (Ahn et al., 2017; Zhang et al., 2020). Given the aim of the current study, we focused on the model parameters relevant to computing social influence (β(N_against_)) and social learning (β(*V*_other_)), respectively (but see **Table 1** for all posterior parameters).

In line with our model-agnostic behavioral results of choice switch probability (**Figure 2A**), the degree each participant weighed dissenting social information (i.e., β(N_against_)) during choice adjustment differed across stimulation sites (one-way repeated-measures ANOVA: F_1,30_ = 9.830, p = 0.004; **Figure 3D**). Further analysis suggested that disruption of the left TPJ decreased the computation of social influence as opposed to both the right TPJ (t_60_ = 4.407, p = 0.0001, Tukey corrected) and the vertex (t_60_ = 2.626, p = 0.029, Tukey corrected), and no significant difference was found between the right TPJ and the vertex (t_60_ = 1.871, p = 0.185, Tukey corrected). Interestingly, no difference was found across stimulation sites regarding how participants integrated vicarious value computation (i.e., β(*V*_other_)) into their own learning processes (one-way repeated measures ANOVA: F_1,30_ = 3.718, p = 0.061; **Figure 3C**), and this result was additionally supported by a Bayes factor (Schmalz et al., 2021) analysis (BF_10_ = 0.106, with default Cauchy priors).

In the last step, we sought to test the association between model parameter and behavior. We reasoned that if the degree of computing social influence was reduced when the left TPJ was stimulated, we ought to anticipate its weaker predictability of switching behavior. Indeed, we found that disrupting the left TPJ led to the weakest effect of β(N_against_) in predicting choice switch probabilities (quantified as slopes computed from **Figure 2A**) with a simple linear regression (rTPJ: b = 0.084, p = 0.004; vertex: b = 0.109, p = 0.031; lTPJ: b = 0.063, p = 0.015; effects compared with Wald tests: p_rTPJ_vertex_ = 0.416, p_rTPJ_lTPJ_ = 0.168, p_lTPJ_vertex_ = 0.003, Bonferroni corrected p-value = 0.05/3 = 0.016; **Figure 3E**). Putting together, our computational modeling effort suggested that disruption of the left TPJ reduced the computation of social influence from the others in the group, whereas the vicarious social learning process by observing the others’ actions remained intact.

## DISCUSSION

Leveraging non-invasive brain stimulation and comprehensive computational modeling, we tested whether left or right TPJ causally supports the computation of social influence in goal-directed learning. We show that disrupting activity in the left TPJ resulted in reduced choice adjustment and declined reaction speed, especially when individuals were contradicted by the entire group. Our model further revealed that disruption of the left TPJ diminished the extent to which social influence was computed, yet the integration of social learning into individuals’ own valuation was intact.

Our previous fMRI results (Zhang and Gläscher, 2020) clear demonstrated that both right and left TPJ encode dissenting social information, but the current results extend these findings such that only left TPJ appears to exert a causal influence on the ensuing behavioral change. Moreover, as we previously reported that bilateral TPJ and the dissenting social information encoded therein in combination with the behavioral adjustment (and the corresponding neural signal in the dorsolateral prefrontal cortex) modulate value computations in prefrontal cortices (ventromedial prefrontal cortex, vmPFC, and anterior cingulate cortex, ACC, encoding direct learning and observational learning, respectively), it is plausible to infer a reduced connectivity pattern between left TPJ and vmPFC/ACC based on the current TMS data. It is worth noting that this reduced connectivity pattern remains speculative and future combined TMS-fMRI experiments will be required.

These findings are arguably in accordance with two lesion studies highlight the role of left TPJ in mentalizing about others: patients with lesions in this part of the brain are impaired in cognitive tasks that require thinking about false beliefs of others and taking their perspective (Samson et al., 2004; Apperly et al., 2007). In addition, a recent fMRI study reported functionally different roles for left and right TPJ in an interactive decision-making task requiring representing about the other’s strategies (Ogawa and Kameda, 2019). They compared human performance against another human participant, an intelligent learning algorithm, and a simple fixed probability rule, and reported that right TPJ showed the largest effect when playing against another human participant, whereas left TPJ exhibited similar effects when playing against a human or an intelligent algorithm. In light of these findings, stimulating right TPJ in our task, in which participants played together with an intelligent agent, might not have led to reduction in social influence, because no other human players were involved. In contrast, the reduction in switch probability and the correspondingly smaller β(N_against_) following TMS of left TPJ might reflect a more general impairment in social reasoning about the others independent of whether the interacting players are human or intelligent artificial algorithms.

A potential caveat of this study revolves around the role of the TPJ in the functions of the dorsal attention network (Corbetta and Shulman, 2002). Stimulating TPJ might have also corrupted attentional selection and the observed effect might simply reflect altered attentional processing rather than the encoding of dissenting social information. However, this is unlikely for several reasons. First, our stimulation site was identified as the overlapping regions between our previous ROIs and the NeuroSynth brain masks resulted from “mentalizing”, rather than “attention”, as the search term. Moreover, a detailed inspection of the brain regions involved in attentional reorientation and ToM reveals spatially separable and only partially overlapping regions in the TPJ (Decety and Lamm, 2007; Carter and Huettel, 2013; Kiesow et al., 2021). Furthermore, an attentional account would leave the differences between left and right TPJ in our task unanswered. Finally, our cognitive modelling revealed that the value signal computed for the other players was non-zero and comparable in all three stimulation conditions making a pure attentional explanation rather unlikely. Nevertheless, in future brain stimulation studies of the TPJ, it would be desirable to measure the representation of social information and attentional processing simultaneously to investigate potential interaction effects more directly.

In conclusion, our study revealed a causal role of left TPJ in encoding dissenting social information about the choices of others and is therefore an essential node in the comprehensive brain network that integrates one’s own and others’ decisions into a coherent value signal that is used for making and updating social decisions.

## MATERIALS AND METHODS

### Data and code availability

All data needed to evaluate the conclusions in the paper are present in the paper and the Supplementary Materials. Behavioral data and custom code to perform analyses can be accessed on the GitHub repository: https://github.com/lei-zhang/SIT_TMS.

### Participants

Forty right-handed participants were invited to participate in the study. No one had any history of neurological and psychiatric diseases, nor currently used medication except contraceptives. Nine participants were excluded for various reasons: one participant had participated in our previous study, one participant had extremely low motor threshold, three participants experienced technical failure (two with experimental program crash and one with TMS device overheat), one participant did not make the choice adjustment, and four participants missed more than 30% of the responses during one or more stimulation. The final sample consisted of 31 participants (17 females). All participants gave informed written consent before the experiment. The study was conducted in accordance with the Declaration of Helsinki and was approved by the Ethics Committee of the Medical Association of Hamburg (PV3661).

### cTBS stimulation sites

We had three stimulation sites in the current experiment, right TPJ, left TPJ, and vertex. We based our bilateral TPJ stimulation sites in Brodmann area 39 on the 2nd-level neuroimaging map from the parametric modulation of dissenting social information from our previous study (Zhang & Gläscher, 2020 Figure S4/Table S4). The peaks (MNI coordinates) of the bilateral TPJ were identified at x = 50, y = –60, z = 34 (right), and x = –48, y = –62, z = 30 (left), respectively, considering the joint areas between our previous 2nd-level map and the meta-analytical maps derived from NeuroSynth (https://neurosynth.org/) using the term “mentalizing” (meta-analysis of 151 studies). Subject-specific stimulation coordinates were obtained using inverse normalization with trilinear interpolation implemented in SPM12 (Statistical Parametric Mapping; Wellcome Trust Center for Neuroimaging, University College London, London, UK) from MNI space to native space. Those coordinates were then superimposed onto each participant’s native structural (T1) images obtained no older than 6 months prior to the experiment. For the non-active control site, we chose the vertex, defined for each participant in their own T1-weighted MRI scan as the intersection of the central sulci from both cerebral hemispheres. Vertex has been commonly used as a control stimulation site as stimulating vertex has minimal task-relevant effects (Polanía et al., 2018). Locating subject-specific stimulation sites, as well as creating landmarks of each participant’s brain, was implemented with the neuronavigation pipeline in the Brainsight software (Rogue Resolutions Inc Montreal, Quebec, Canada). All participants were blinded as to the stimulation sites and the neuronavigation setup.

### Stimulation protocols

We applied a cTBS transcranial magnetic stimulation (TMS) protocol to each participant-specific coordinates identified with the above procedure, with the handle pointing posteriorly. Following previous cTBS studies on social neuroscience (e.g., Hill et al., 2017; Bukowski et al., 2020), the cTBS stimulation protocol comprised 600 pulses administered for 40 s, in bursts of 3 pulses at 50 Hz (20 ms) repeated at intervals of 5 Hz (200 ms). Stimulations were controlled and delivered using the Magstim-Rapid-2 stimulator with an air-cooled 70 mm figure-8 coil (Magstim Co Ltd. Spring Gardens, Whitland, UK). The stimulation intensity was determined as 80% of the active motor threshold. Motor threshold was determined as the lowest single TMS pulse intensity required (through a staircase procedure) to elicit a slightly visible twitch of the thumb and/or the index finger on in more than 5 out of 10 times stimulation while participants maintained a constant pressure between the thumb and the index finger at 20% of maximum force. We employed a within-subject design across three stimulation sites. To prevent potential carry over effect, the stimulation of the vertex was always kept in the middle, while the order between the right and the left TPJ was counterbalanced. Each experimental session lasted shorter than 25 minutes, which was adequately within the hypothesized duration of disrupted excitability at the stimulated area (Romero et al., 2020).

### Experimental task

The core of our social learning task was a probabilistic reversal learning (PRL) paradigm, where each choice option was associated with a particular reward probability (i.e., 70% and 30%). After a variable length of trials (i.e., 8-12 trials), the reward contingencies reversed, such that individuals needed to re-adapt to the new reward contingencies in order to maximize their outcome. That said, the PRL task assured constant learning throughout the entire experiment.

Different from our previous study involving real-time interactions of 5 participants (Zhang and Gläscher, 2020), in the current study only one participant was tested. For each experimental session, participants were informed that they were about to play with four independent “intelligent computer algorithms” that best matched human behavior in the previous study. Hence, there was no deception in the current study. Importantly, participants were instructed that those computer algorithms were able to learn from their own errors, and also take decisions of the others into consideration (i.e., the other three algorithms together with the participant). In fact, these algorithms were simulated from the best computational model in our full version social learning task (Zhang and Gläscher, 2020). To better maintain participants’ attention and ecological validity, we used human faces to indicate the computer algorithms during the experimental presentation.

The task consisted of 3 phases for every trial. (i) Phase 1. Initial choice (Choice 1). Upon the presentation of two choice options using abstract fractals, participants were asked to make their 1st choice (2800 ms). A yellow frame was then presented to highlight the chosen option. (ii) Phase 2. Choice adjustment (Choice 2). When all four other choices were presented, participants were able to adjust their choices given the social information (2800 ms). The yellow frame was shifted accordingly to highlight the adjusted choice. (iii) Phase 3. Outcome delivery. Finally, the outcome was determined by participants’ 2nd choice (3000 ms plus a jittered inter-trial interval 2000 – 4000 ms; Figure 1A). Outcomes of the other four players were also displayed. On each trial, the reward was assigned to only one choice option given the reward probability, whereas choosing the other option would lead to a punishment. The reward realization sequence (trial-by-trial complementary win and loss) was generated with a pseudo-random order. Three pairs of abstract fractals were used in the current study, with a fully counterbalanced assignment together with the stimulation sites. All participants were compensated with a base payment of 20 Euro plus the average reward they achieved across three sessions of the experiment. Finally, the experiment ended with an informal debriefing session.

### Behavioral data acquisition

Stimulus presentation and response recording were accomplished with Matlab R2014b (www.mathworks.com) and Cogent2000 (www.vislab.ucl.ac.uk/cogent.php). Buttons of “V” and “B” on the keyboard corresponded to the left and right choice options, respectively. To avoid motor artifacts, the position of the two choices options was counterbalanced for all participants.

### Behavioral data analysis

We tested the cTBS effects on participants’ choice adjustment after observing the social information (during Phase 3 of the task), by three key measurements: (1) choice switch probability, (2) response time (RT), and (3) choice accuracy. According to our previous finding, here, we were particularly interested in the stimulation effect on the interaction between the relative direction of the group (with vs. against) and group consensus (2:2, 3:1, and 4:0, view of each participant). Therefore, we computed the difference measurement between the “against” and “with” conditions at each group consensus level as the dependent variable of interest. Accordingly, we performed 3 (cTBS sites: right TPJ, vertex, left TPJ) × 3 (group consensus: 2:2, 3:1, and 4:0) linear mixed models (LMM), with stimulation sites and group consensus as fixed factors (with interaction) and random slopes (without interaction), and participants as random intercept: y ∼ stimulation * consensus + (1 + stimulation + consensus | ID). This LMM structure was identical for the analyses of Choice 2’s switch probability, RT, and accuracy.

All statistical tests were performed in R (v3.7.1; www.r-project.org). All repeated-measures LME models were analyzed with the “lme4” package and summarized with the “BruceR” package (https://github.com/psychbruce/bruceR). Results were considered statistically significant at the level *p* < 0.05. Multiple comparison correction was applied whenever appropriate.

### Computational modeling

The computational modeling procedures are fully documented in our previous work (Zhang and Gläscher, 2020), and for the consideration of enhancing accessibility and reproducibility, we repeat the main points relevant to the current study herein. Note that, we deem that it is the best practice to deviate as little as possible from our previous modeling description.

We constructed three categories of models: baseline non-social model (M1), social model (M2) with only social influence (before receiving the outcome), and social model (M3–M6) with both social influence and social learning (before receiving the outcome).

In all models, Choice 1 was accounted for by the option values of option A and option B:

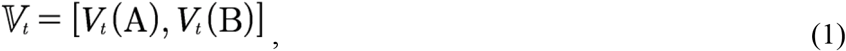

where *𝕍*_*t*_ indicated a two-element vector consisting of option values of A and B on trial *t*. Values were then converted into action probabilities using a Softmax function. On trial *t*, the action probability of choosing option A was defined as follows:

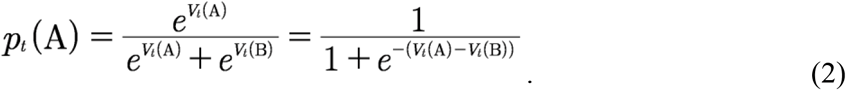

For Choice 2, we modeled it as a “switch” (coded as 1) or a “stay” (coded as 0) with respect to Choice 1 using a logistic regression. On trial *t*, the probability of switching given the switch value was defined as follows:

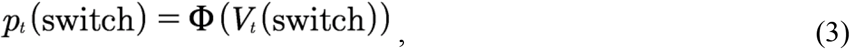

where Φ was the inverse logit linking function:

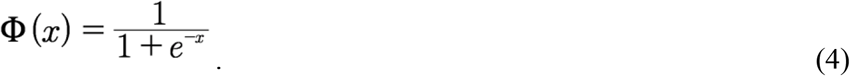

It is worth noting that, in model specifications of the action probability, we did not include the commonly used inverse Softmax temperature parameter *τ* (except M1). This was because we explicitly constructed the option values of Choice 1 and the switch value of Choice 2 in a design-matrix fashion. Therefore, including the inverse Softmax temperature parameter would inevitably give rise to a multiplication term, which, as a consequence, would cause unidentifiable parameter estimation.

In the simplest model (M1), a fictitious update Rescorla-Wagner model was used to model the Choice 1, as we have demonstrated that the fictitious update model outperformed the standard single update Rescorla-Wagner (Rescorla and Wagner, 1972) model and Pearce-Hall (Pearce and Hall, 1980) model with dynamic learning rate. Here, both the chosen value and the unchosen value were updated, as in:

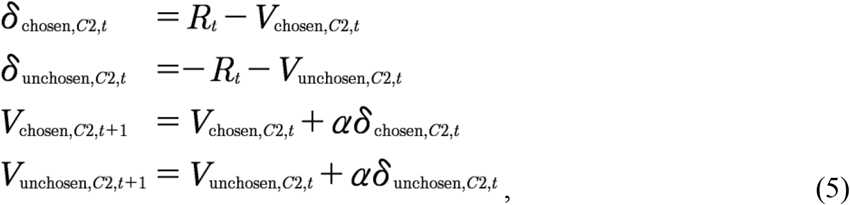

where *R*_*t*_ was the outcome on trial *t*, and *α* (0 < *α* < 1) denoted the learning rate that accounted for the weight of reward prediction error in value update. A beta weight (*β*_V_; akin to the inverse temperature parameter) was multiplied with the values before being submitted to Eq. 2 with a Categorical distribution, as in:

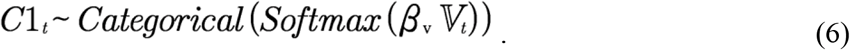

Because there was no social information in M1a, the switch value of Choice 2 was comprised merely of the value difference of Choice 1 and a switching bias (i.e., intercept):

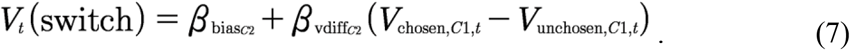

Choice 2 was then modeled with this switch value following a Bernoulli distribution:

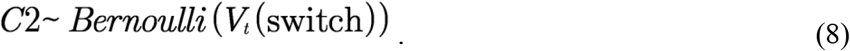

M2 tested whether observing choices from the other players (i.e., social influence) contributed to the choice switching. As on top of M1, only the switch value of Choice 2 was modified:

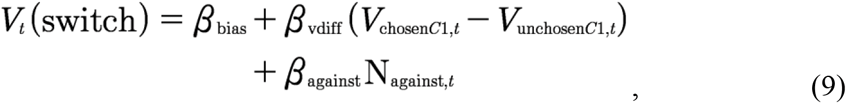

where N_against,*t*_ denoted the amount of dissenting social information relative to each participant’s Choice 1 on trial *t*.

M3–M6 assessed whether participants learned from their social partners and whether they updated vicarious option values through social learning, by testing several competing hypotheses of how vicarious valuation contributed to Choice 1 on the following trial. Across M3–M6, the option values of Choice 1 was specified by a weighted combination between *V*_self_ updated via direct learning and *V*_other_ updated via social learning, and M3–M6 differed on the specification of *V*_other_.

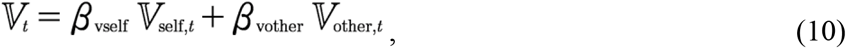

where

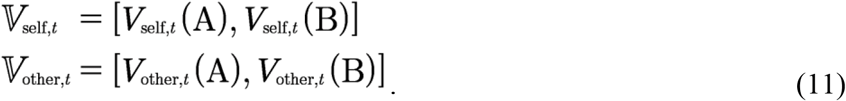

M3 tested whether individuals recruited a similar RL algorithm to their own when learning option values from observing others. As such, M3 assumed participants to update values “for” the others using the same fictitious update rule for themselves:

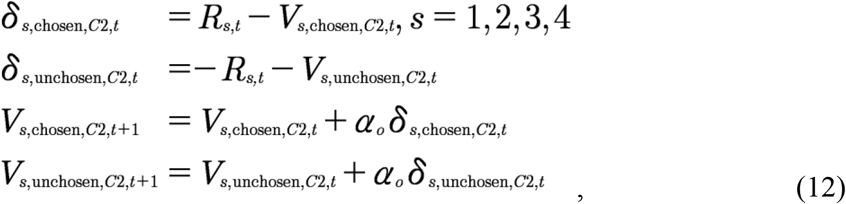

where *s* denoted the index of the four other co-players. These option values from the four co-players were then preference-weighted and summed to formulate *V*_other_, as follows:

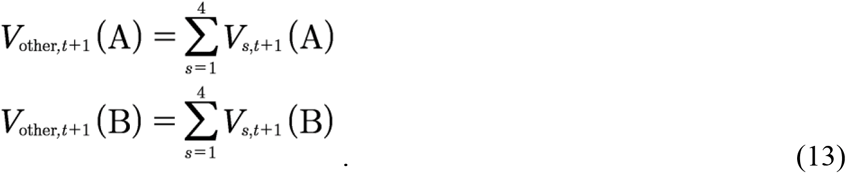

To ensure that the corresponding value-related parameters (*β*_Vself_ and *β*_Vother_ in Eq. 10) were comparable, *V*_other_ (across M3–M6) was further normalized to lie between –1 and 1 with the Φ(x) function defined in Eq. 4:

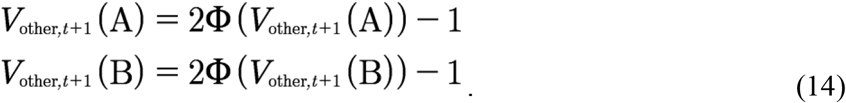

One may argue that having four independent RL agents as in M3 was cognitively demanding. We, therefore, constructed three additional models (M4–M6) that employed simpler but distinct computations to update vicarious values via social learning. Now that M3 considered both choice and outcome to determine the action value, we asked if using either choice or outcome alone may perform as well as, or even better than, M3. That said, we constructed M4 that updated *V*_other_ using only the others’ action preference, M5 that considered the others’ current outcome, and M6 that tracked the others’ cumulative outcome, to resemble the value update via observational learning.

In M4, other players’ action preference (*ρ*) is derived from the choice history over the last three trials (from T-2 to T) using the cumulative distribution function of the beta distribution at the value of 0.5 (*I*_0.5_). That is:

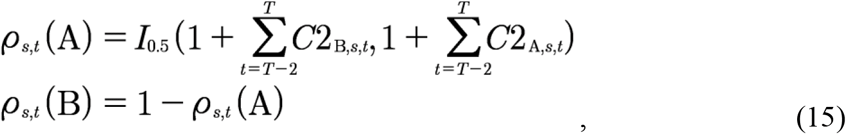

To illustrate, if one co-player chose option A twice and option B once, the action preference of choosing A for him/her was: *I*_0.5_(frequency of B + 1, frequency of A + 1) = *I*_0.5_(0.5, 1 + 1, 2 + 1) = 0.6875. *V*_other_ was computed based on these action preferences:

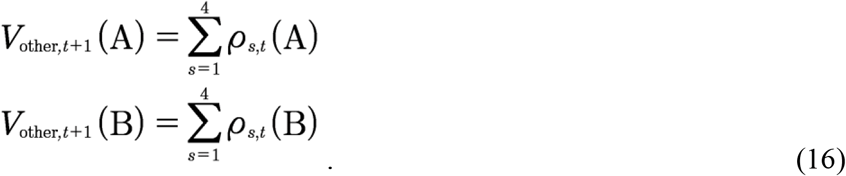

By contrast, M5 tested whether participants updated *V*_other_ using only each other’s reward on the current trial, which was equivalent to the standard Rescorla-Wagner model with *α* = 1, indicating no trial-by-trial learning:

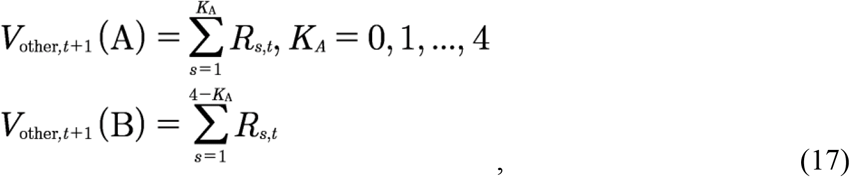

where *K*_A_ denoted the number of other players who decided on option A on trial *t*.

Lastly, M6 assessed whether participants tracked cumulated reward histories over the last few trials instead of monitoring only the most recent outcome of the others, with a discounted reward history over the last three trials:

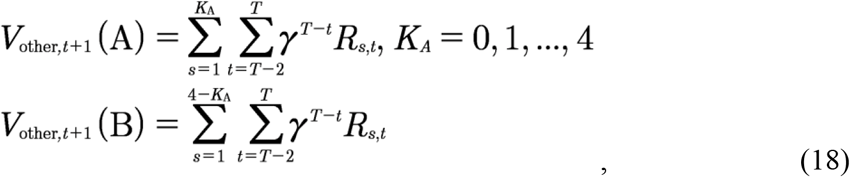

where *γ* (0 < *γ* < 1) denoted the rate of exponential decay, all other notions were as in Eq. 17.

### Model estimation with hierarchical Bayesian analysis

Model estimations of all candidate models were performed with hierarchical Bayesian analysis (HBA; Gelman et al., 2013) using a newly developed statistical computing language Stan (Carpenter et al., 2017) in R, following the implementation in the “hBayesDM” package (Ahn et al., 2013). Stan utilizes a Hamiltonian Monte Carlo (HMC; and efficient Markov Chain Monte Carlo, MCMC) sampling scheme to perform full Bayesian inference and obtain the actual posterior distribution. We additionally implemented effect coding to account inter-dependencies in within-subject experimental design. Let φ_*s,c*_ denote a generic individual-level parameter of participant *s* in stimulation site *c*. φ_*s,c*_ was drawn from a group-level multivariate normal distribution:

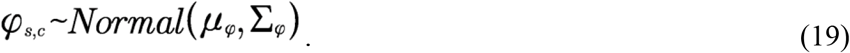

Here, μ_*φ*_ was a three-element vector (1: vertex, 2: right TPJ, 3: left TPJ) of group-level means, where μ_2_, and μ_3_ were effect-coded as the difference with respect to μ_1_:

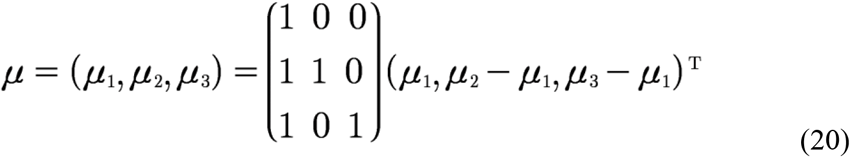

Σ_φ_ was a variance-covariance matrix denoting the group-level multivariate standard deviation, and it could be decomposed with the corresponding correlation matrix Ω_φ_ of correlation coefficients *r*:

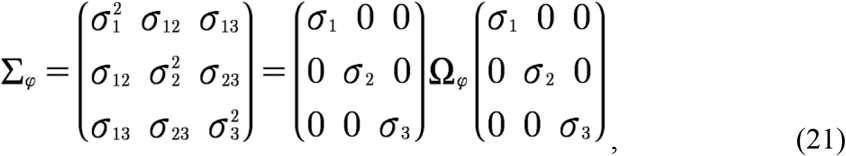

where,

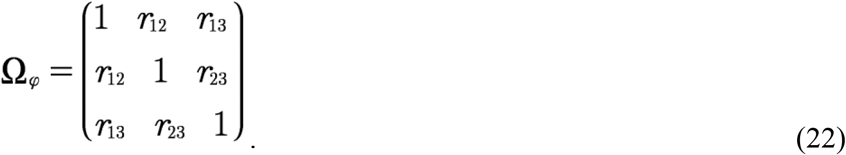

And lastly, the correlation matrix was reparameterized by Cholesky decomposition factor, noted as two triangular matrices (*L*_*φ*_*L*_*φ*_’). All these group-level parameters were specified with weakly-informative priors (Gelman et al., 2013): μ_φ_ ∼ Normal (0, 1), Σ_φ_.∼ half-Cauchy (0, 3), and *L*_φ_ ∼ LJK (2). All parameters were unconstrained except for α and γ (both [0 1] constraint, with inverse probit transform). We fit each candidate model with four independent MCMC chains using 1000 iterations after 1000 iterations for the initial algorithm warmup per chain, which resulted in 4000 valid posterior samples. The convergence of the MCMC chains was assessed both visually (from the trace plot) and through the Gelman-Rubin 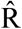 Statistics (Gelman and Rubin, 1992). 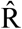 values of all parameters were close to 1.0 (at most smaller than 1.05 in the current study), which indicated adequate convergence.

For model comparison, we computed the Leave-One-Out information criterion (LOOIC) score per candidate model (Vehtari et al., 2016). The LOOIC score provides the point-wise estimate (using the entire posterior distribution) of out-of-sample predictive accuracy in a fully Bayesian way, which is more reliable compared to point-estimate information criterion (e.g., Akaike information criterion, AIC). We additionally performed Bayesian model averaging (BMA) with Bayesian bootstrap (Yao et al., 2018) to compute the probability of each candidate model being the best model.

## SUPPLEMENTARY INFORMATION

Supplemental Information includes 3 tables can be found with this article at http://xxxx.

## ACKNOWLEDGMENTS

We thank Vivien Breckwoldt and Kevin Rozario for help with data acquisition. L.Z. was supported the Research Promotion Fund (FFM) for young scientists of the University Medical Center Hamburg-Eppendorf, the Vienna Science and Technology Fund (WWTF VRG13-007), and the Austrian Science Fund (FWF-M3166). J.G., C.C.H., F.I.K, K.Z., and X.F. were supported by the Collaborative Research Centers “Cross-modal learning” (DFG TRR 169). C.C.H. was additionally supported by the Landesforschungsförderung FHH, CRC 936 “Multisite Communication in the Brain”, and the Human Brain Project HBP SGA3. Furthermore, J.G. was supported by the Bernstein Award for Computational Neuroscience (BMBF 01GQ1006), the Collaborative Research Center “Cognition of Interaction” (DFG SFB 1528), and a Collaborative Research in Computational Neuroscience (CRCNS) grant (BMBF 01GQ1603).

## AUTHOR CONTRIBUTIONS

**L.Z**.: Conceptualization, Data curation, Formal Analysis, Funding acquisition, Investigation, Methodology, Project administration, Data Acquisition, Software, Validation, Visualization, Writing – original draft, Writing – review & editing. **F.I.K**.: Investigation, Methodology, Resources, Validation, Writing – review & editing. **K.Z**.: Validation, Writing – review & editing. **X.F**.: Validation, Writing – review & editing. **C.L**.: Validation, Writing – review & editing. **C.C.H**.: Funding acquisition, Resources, Supervision, Validation, Writing – review & editing. **J.G**.: Conceptualization, Formal Analysis, Funding acquisition, Methodology, Data Acquisition, Project administration, Resources, Supervision, Validation, Writing – original draft, Writing – review & editing.

## REFERENCES

Ahn W-Y, Haines N, Zhang L. 2017. Revealing neurocomputational mechanisms of reinforcement learning and decision-making with the hBayesDM package. Comput Psychiatry 1:24–57. doi:10.1162/CPSY_a_00002

Apperly IA, Samson D, Chiavarino C, Bickerton W-L, Humphreys GW. 2007. Testing the domain-specificity of a theory of mind deficit in brain-injured patients: Evidence for consistent performance on non-verbal, “reality-unknown” false belief and false photograph tasks. Cognition 103:300–321. doi:10.1016/j.cognition.2006.04.012

Baumgartner T, Schiller B, Rieskamp J, Gianotti LRR, Knoch D. 2014. Diminishing parochialism in intergroup conflict by disrupting the right temporo-parietal junction. Soc Cogn Affect Neurosci 9:653– 660. doi:10.1093/scan/nst023

Bukowski H, Tik M, Silani G, Ruff CC, Windischberger C, Lamm C. 2020. When differences matter: rTMS/fMRI reveals how differences in dispositional empathy translate to distinct neural underpinnings of self-other distinction in empathy. Cortex 128:143–161. doi:10.1016/j.cortex.2020.03.009

Carpenter B, Gelman A, Hoffman MD, Lee D, Goodrich B, Betancourt M, Brubaker M, Guo J, Li P, Riddell A. 2017. Stan : A Probabilistic Programming Language. J Stat Softw 76:1–32. doi:10.18637/jss.v076.i01

Carter RM, Huettel SA. 2013. A nexus model of the temporal–parietal junction. Trends Cogn Sci 17:328– 336. doi:10.1016/j.tics.2013.05.007

Charness G, Gneezy U, Kuhn MA. 2012. Experimental methods: Between-subject and within-subject design. J Econ Behav Organ 81:1–8. doi:10.1016/j.jebo.2011.08.009

Charpentier CJ, Iigaya K, O’Doherty JP. 2020. A Neuro-computational Account of Arbitration between Choice Imitation and Goal Emulation during Human Observational Learning. Neuron 106:687–699.e7. doi:10.1016/j.neuron.2020.02.028

Corbetta M, Shulman GL. 2002. Control of goal-directed and stimulus-driven attention in the brain. Nat Rev Neurosci 3:201–215. doi:10.1038/nrn755

Decety J, Lamm C. 2007. The role of the right temporoparietal junction in social interaction: How low-level computational processes contribute to meta-cognition. Neuroscientist 13:580–593. doi:10.1177/1073858407304654

Deschrijver E, Palmer C. 2020. Reframing social cognition: Relational versus representational mentalizing. Psychol Bull 146:941–969. doi:10.1037/bul0000302

Gelman A, Rubin DB. 1992. Inference from iterative simulation using multiple sequences. Stat Sci 7:457–472. doi:10.1214/ss/1177011136

Gelman A, Stern HS, Carlin JB, Dunson DB, Vehtari A, Rubin DB. 2013. Bayesian data analysis. Boca Raton, FL: Chapman and Hall/CRC.

Gläscher J, Hampton AN, O’Doherty JP. 2009. Determining a role for ventromedial prefrontal cortex in encoding action-based value signals during reward-related decision making. Cereb Cortex 19:483– 495. doi:10.1093/cercor/bhn098

Hill C, Suzuki S, Polanía R, Moisa M, O’Doherty JP, Ruff CC. 2017. A causal account of the brain network computations underlying strategic social behavior. Nat Neurosci 20:1142–1149. doi:10.1038/nn.4602

Huang Y-Z, Edwards MJ, Rounis E, Bhatia KP, Rothwell JC. 2005. Theta Burst Stimulation of the Human Motor Cortex. Neuron 45:201–206. doi:10.1016/j.neuron.2004.12.033

Kiesow H, Spreng RN, Holmes AJ, Chakravarty MM, Marquand AF, Yeo BTT, Bzdok D. 2021. Deep learning identifies partially overlapping subnetworks in the human social brain. Commun Biol 4:65. doi:10.1038/s42003-020-01559-z

Konovalov A, Hill C, Daunizeau J, Ruff CC. 2021. Dissecting functional contributions of the social brain to strategic behavior. Neuron 109:3323–3337.e5. doi:10.1016/j.neuron.2021.07.025

Obeso I, Moisa M, Ruff CC, Dreher J-C. 2018. A causal role for right temporo-parietal junction in signaling moral conflict. Elife 7:1–16. doi:10.7554/eLife.40671

Ogawa A, Kameda T. 2019. Dissociable roles of left and right temporoparietal junction in strategic competitive interaction. Soc Cogn Affect Neurosci 14:1037–1048. doi:10.1093/scan/nsz082

Pearce JM, Hall G. 1980. A model for Pavlovian learning: Variations in the effectiveness of conditioned but not of unconditioned stimuli. Psychol Rev 87:532–552. doi:10.1037/0033-295X.87.6.532

Polanía R, Nitsche MA, Ruff CC. 2018. Studying and modifying brain function with non-invasive brain stimulation. Nat Neurosci 21:174–187. doi:10.1038/s41593-017-0054-4

Rescorla RA, Wagner AR. 1972. A theory of Pavlovian conditioning: Variations in the effectiveness of reinforcement and nonreinforcementClassical Conditioning II: Current Research and Theory. New York, NY: Appleton-Century-Crofts. pp. 64–99.

Romero MC, Merken L, Janssen P, Davare M. 2020. Neural effects of continuous theta-burst stimulation in macaque parietal neurons. bioRxiv. doi:10.1101/2020.12.07.414482

Ruff CC, Fehr E. 2014. The neurobiology of rewards and values in social decision making. Nat Rev Neurosci 15:549–562. doi:10.1038/nrn3776

Rusch T, Steixner-Kumar S, Doshi P, Spezio M, Gläscher J. 2020. Theory of mind and decision science: Towards a typology of tasks and computational models. Neuropsychologia 146:107488. doi:10.1016/j.neuropsychologia.2020.107488

Samson D, Apperly IA, Chiavarino C, Humphreys GW. 2004. Left temporoparietal junction is necessary for representing someone else’s belief. Nat Neurosci 7:499–500. doi:10.1038/nn1223

Schaafsma SM, Pfaff DW, Spunt RP, Adolphs R. 2015. Deconstructing and reconstructing theory of mind. Trends Cogn Sci 19:65–72. doi:10.1016/j.tics.2014.11.007

Schmalz X, Biurrun Manresa J, Zhang L. 2021. What is a Bayes factor? Psychol Methods. doi:10.1037/met0000421

Schurz M, Radua J, Tholen MG, Maliske L, Margulies DS, Mars RB, Sallet J, Kanske P. 2021. Toward a hierarchical model of social cognition: A neuroimaging meta-analysis and integrative review of empathy and theory of mind. Psychol Bull 147:293–327. doi:10.1037/bul0000303

Vehtari A, Gelman A, Gabry J. 2016. Practical Bayesian model evaluation using leave-one-out cross-validation and WAIC. Stat Comput 27:1–20. doi:10.1007/s11222-016-9696-4

Yao Y, Vehtari A, Simpson D, Gelman A. 2018. Using Stacking to Average Bayesian Predictive Distributions (with Discussion). Bayesian Anal 13:917–1007. doi:10.1214/17-BA1091

Yarkoni T, Poldrack RA, Nichols TE, Van Essen DC, Wager TD. 2011. Large-scale automated synthesis of human functional neuroimaging data. Nat Methods 8:665–670. doi:10.1038/nmeth.1635

Zhang L, Gläscher J. 2020. A brain network supporting social influences in human decision-making. Sci Adv 6:eabb4159. doi:10.1126/sciadv.abb4159

Zhang L, Lengersdorff L, Mikus N, Gläscher J, Lamm C. 2020. Using reinforcement learning models in social neuroscience: frameworks, pitfalls and suggestions of best practices. Soc Cogn Affect Neurosci 15:695–707. doi:10.1093/scan/nsaa089

