## Supplemental Information for "A causal role of the human left temporoparietal junction in computing social influence during goal-directed learning"

**Supplemental Figures**

**Supplemental Tables**

| ANOVA Table | | | | | |
|---|---|---|---|---|---|
| | SumSq | NumDF | DenDF | *F* | *p* |
| stim | 0.06 | 2 | 30.07 | 1.40 | 0.262 |
| cons | 0.47 | 2 | 51.69 | 11.46 | < 0.001 |
| stim:cons | 0.32 | 4 | 179.98 | 3.90 | 0.005 |

| Beta Table | | | | | |
|---|---|---|---|---|---|
| | Beta | 95% CI | SE | *t* | *p* |
| stim(rtpj) | 0 | [-0.179, 0.179] | 0.090 | 0 | 1 |
| stim(ltpj) | 0 | [-0.160, 0.160] | 0.081 | 0 | 1 |
| cons(3:1) | 0.241 | [ 0.069, 0.413] | 0.087 | 2.77 | 0.007 |
| cons(4:0) | 0.367 | [ 0.159, 0.575] | 0.104 | 3.53 | < 0.001 |
| stim(rtpj):cons(3:1) | 0.044 | [-0.094, 0.182] | 0.070 | 0.62 | 0.534 |
| stim(ltpj):cons(3:1) | 0.014 | [-0.125, 0.152] | 0.070 | 0.19 | 0.847 |
| stim(rtpj):cons(4:0) | 0.155 | [ 0.017, 0.293] | 0.070 | 2.22 | 0.028 |
| stim(ltpj):cons(4:0) | -0.095 | [-0.234, 0.043] | 0.070 | -1.36 | 0.174 |

**Table S1.**

**Linear mixed-effect model results for the measurement of choice switch probability, related to Figure 2A and STAR methods**.

ANOVA table shows type III statistics of analysis of variance (ANOVA). Sum Sq, sum of squares; NumDF, numerator degrees of freedom; DenDF, denominator degrees of freedom (estimated by Satterthwaite approximation); *F*, *F*-statistics; *p*, *p*-value. Stim, stimulation sites (i.e., rTPJ, lTPJ, vertex); cons, group consensus (i.e., 2:2, 3:1, 4:0); stim:cons, interaction between stimulation sites and group consensus. Beta table shows estimates of coefficients. Beta, standardized coefficients; 95% CI, 95% confidence interval of beta coefficients; SE, standard error of coefficients; *t*, *t*-statistics.

| ANOVA Table | | | | | |
|---|---|---|---|---|---|
| | SumSq | NumDF | DenDF | *F* | *p* |
| stim | 50.007 | 2 | 40.56 | 1.44 | 0.249 |
| cons | 4.149 | 2 | 34.88 | 0.12 | 0.888 |
| stim:cons | 91.306 | 4 | 180.00 | 1.31 | 0.267 |

| Beta Table | | | | | |
|---|---|---|---|---|---|
| | Beta | 95% CI | SE | *t* | *p* |
| stim(rtpj) | 0 | [-0.187, 0.187] | 0.095 | 0 | 1 |
| stim(ltpj) | 0 | [-0.229, 0.229] | 0.115 | 0 | 1 |
| cons(3:1) | -0.033 | [-0.233, 0.167] | 0.101 | -0.33 | 0.745 |
| cons(4:0) | -0.087 | [-0.312, 0.138] | 0.113 | -0.77 | 0.444 |
| stim(rtpj):cons(3:1) | 0.006 | [-0.170, 0.182] | 0.089 | 0.07 | 0.948 |
| stim(ltpj):cons(3:1) | 0.093 | [-0.083, 0.269] | 0.089 | 1.04 | 0.298 |
| stim(rtpj):cons(4:0) | 0.058 | [-0.118, 0.234] | 0.089 | 0.65 | 0.518 |
| stim(ltpj):cons(4:0) | 0.197 | [ 0.021, 0.373] | 0.089 | 2.21 | 0.028 |

**Table S2.**

**Linear mixed-effect model results for the measurement of response time of Choice 2, related to Figure 2B and STAR methods**.

ANOVA table shows type III statistics of analysis of variance (ANOVA). Sum Sq, sum of squares; NumDF, numerator degrees of freedom; DenDF, denominator degrees of freedom (estimated by Satterthwaite approximation); *F*, *F*-statistics; *p*, *p*-value. Stim, stimulation sites (i.e., rTPJ, lTPJ, vertex); cons, group consensus (i.e., 2:2, 3:1, 4:0); stim:cons, interaction between stimulation sites and group consensus. Beta table shows estimates of coefficients. Beta, standardized coefficients; 95% CI, 95% confidence interval of beta coefficients; SE, standard error of coefficients; *t*, *t*-statistics.

| ANOVA Table | | | | | |
|---|---|---|---|---|---|
| | SumSq | NumDF | DenDF | *F* | *p* |
| stim | 0.09 | 2 | 34.48 | 0.49 | 0.617 |
| cons | 0.90 | 2 | 130.07 | 4.66 | 0.011 |
| stim:cons | 0.20 | 4 | 130.00 | 0.52 | 0.719 |

| Beta Table | | | | | |
|---|---|---|---|---|---|
| | Beta | 95% CI | SE | *t* | *p* |
| stim(rtpj) | 0.008 | [-0.238, 0.254] | 0.124 | 0.07 | 0.947 |
| stim(ltpj) | 0.041 | [-0.207, 0.289] | 0.125 | 0.33 | 0.744 |
| cons(3:1) | 0.326 | [ 0.053, 0.599] | 0.138 | 2.36 | 0.020 |
| cons(4:0) | 0.205 | [-0.051, 0.460] | 0.129 | 1.59 | 0.115 |
| stim(rtpj):cons(3:1) | -0.104 | [-0.364, 0.156] | 0.132 | -0.79 | 0.431 |
| stim(ltpj):cons(3:1) | -0.093 | [-0.350, 0.165] | 0.130 | -0.71 | 0.476 |
| stim(rtpj):cons(4:0) | -0.028 | [-0.256, 0.199] | 0.115 | -0.25 | 0.806 |
| stim(ltpj):cons(4:0) | -0.137 | [-0.371, 0.097] | 0.118 | -1.16 | 0.247 |

**Table S3.**

**Linear mixed-effect model results for the measurement of accuracy of Choice 2, related to Figure 2C and STAR methods**.

ANOVA table shows type III statistics of analysis of variance (ANOVA). Sum Sq, sum of squares; NumDF, numerator degrees of freedom; DenDF, denominator degrees of freedom (estimated by Satterthwaite approximation); *F*, *F*-statistics; *p*, *p*-value. Stim, stimulation sites (i.e., rTPJ, lTPJ, vertex); cons, group consensus (i.e., 2:2, 3:1, 4:0); stim:cons, interaction between stimulation sites and group consensus. Beta table shows estimates of coefficients. Beta, standardized coefficients; 95% CI, 95% confidence interval of beta coefficients; SE, standard error of coefficients; *t*, *t*-statistics.
